# Accurate assembly of minority viral haplotypes from next-generation sequencing through efficient noise reduction

**DOI:** 10.1101/264242

**Authors:** Sergey Knyazev, Viachaslau Tsyvina, Anupama Shankar, Andrew Melnyk, Alexander Artyomenko, Tatiana Malygina, Yuri B. Porozov, Ellsworth M. Campbell, Serghei Mangul, William M. Switzer, Pavel Skums, Alex Zelikovsky

## Abstract

Rapidly evolving RNA viruses continuously produce minority haplotypes that can become dominant if they are drug-resistant or can better evade the immune system. Therefore, early detection and identification of minority viral haplotypes may help to promptly adjust the patient's treatment plan preventing potential disease complications. Minority haplotypes can be identified using next-generation sequencing (NGS), but sequencing noise hinders accurate identification. The elimination of sequencing noise is a non-trivial task that still remains open. Here we propose CliqueSNV based on extracting pairs of statistically linked mutations from noisy reads. This effectively reduces sequencing noise and enables identifying minority haplotypes with the frequency below the sequencing error rate. We comparatively assess the performance of CliqueSNV using an *in vitro* mixture of nine haplotypes that were derived from the mutation profile of an existing HIV patient. We show that CliqueSNV can accurately assemble viral haplotypes with frequencies as low as 0.1% and maintains consistent performance across short and long bases sequencing platforms.

## INTRODUCTION

Rapidly evolving RNA viruses such as human immunodeficiency virus (HIV), hepatitis C virus (HCV), influenza A virus (IAV), SARS, and SARS-CoV-2 form populations of closely related genomic variants inside infected hosts (1, 2, 3, 4, 5, 6, 7, 8, 9, 10). The intra-host viral populations include minority viral variants that are frequently responsible for drug resistance, immune escape, and disease transmission (11,12,13,14,15,16,17,18,19,20,21,22,23). Therefore, accurately predicting minority viral populations from extremely large and noisy viral genomic data is important for biomedical research, epidemiology, and clinical applications. Although this problem has recently attracted significant interest from the biomedical research community (24,25,26), numerous obstacles still delay NGS integration into the viral studies. The last decade witnessed numerous attempts to employ NGS and bioinformatics methods for reconstructing intra-host viral populations. These methods are not accurate enough for clinical and epidemiological applications since they cannot reliably identify haplotypes accounting for a substantial portion of the population. Existing methods are ill-equipped to assemble closely related haplotypes and have elevated false-positive rates. Additionally, there is only one in vitro viral sequencing benchmark for validation of haplotyping tools (26), and to convincingly demonstrate that such tools are ready for clinical and epidemiological applications, new comprehensive sequencing benchmarks are urgently required (27).

Next-generation sequencing (NGS) technologies now provide versatile opportunities to study viral populations. In particular, the popular Illumina MiSeq/HiSeq platforms produce 25-320 million reads, which allow multiple coverage of highly variable viral genomic regions. This high coverage is essential for capturing rare variants. Ability of NGS technologies to efficiently identify minority variants have recently gained FDA approval (28). However, *haplotyping* of heterogeneous viral populations (i.e., assembly of full-length genomic variants and estimation of their frequencies) is extremely complicated due to the vast number of sequencing reads, the need to assemble an unknown number of closely related viral sequences and to identify and preserve low-frequency variants. Single-molecule sequencing technologies, such as PacBio, provide an alternative to short-read sequencing by allowing full-length viral variants to be sequenced in a single pass. However, the high level of sequence noise due to background or platform-specific sequencing errors produced by all currently available platforms makes inference of low-frequency genetically close variants especially challenging, since it is required to distinguish between real and artificial genetic heterogeneity produced by sequencing errors.

Recently, a number of computational tools for inference of viral quasispecies populations from NGS reads have been proposed (27), including Savage (24), PredictHaplo (29), aBayesQR (30), QuasiRecomb (31), HaploClique (32), VGA (33), VirA (34, 35), SHORAH (36), ViSpA (37), QURE (38) and others (39, 40, 41, 42, 43). Even though these algorithms proved useful in many applications, accurate and scalable viral haplotyping remains a challenge. In particular, inference of low-frequency viral variants is still problematic, while many computational tools designed for the previous generation of sequencing platforms have severe scalability problems when applied to datasets produced by state-of-the-art technologies.

Previously, several tools such as V-phaser (44), V-phaser2 (45) and CoVaMa (46) exploited linkage of mutations for single nucleotide variant (SNV) calling rather than haplotype assembly, but they do not accommodate sequencing errors when deciding whether two variants are linked. These tools are also unable to detect the frequency of mutations above sequencing error rates (47). The 2SNV algorithm (48) accommodates errors in links and was the first such tool to be able to correctly detect haplotypes with a frequency below the sequencing error rate.

We propose a novel method that can accurately identify minority haplotypes from NGS reads consisting of three steps. First, we extract pairs of statistically linked mutations. Second, we find maximal sets of pairwise linked mutations (cliques) where each clique corresponds to a set of mutations in a minority haplotype. Finally, we assign each read to the closest clique, and for each clique, we form a haplotype as a consensus of reads assigned to it.

All haplotyping tools require solid and convincing validation benchmarks (49, 50). The true viral variants and their distribution are only known for simulated data (51), but sequencing errors, variation of coverage depth, PCR bias, and systematic noise cannot be reliably simulated. Therefore experimental sequencing benchmarks that provide an adequate evaluation of haplotyping tools are necessary.

By now, there are only two experimental sequencing benchmarks -(i) Illumina sequencing reads consisting of a mixture of five HIV-1 strains (HIV5exp, see Table 1) (52) and (ii) PacBio sequencing reads from a sample consisting of ten IAV viral variants (IAV10exp, see Table 1) (48). In the HIV5exp, five different HIV-1 strains each having 20% frequency were prepared to mimic an intra-host viral population. Unfortunately, this benchmark is not realistic enough since the observed intra-host viral populations consist of variants that are much closer to each other than different strains and contain both frequent and rare variants (53). The IAV10exp benchmark significantly better mimics the intrahost viral population since its variants are very similar to each other and the variant frequencies are realistically non-uniform. Thus, similar to the IAV10exp benchmark, it would be beneficial to develop Illumina benchmarks which adequately imitate intra-host viral populations containing closely related minority variants.

**Table 1.**
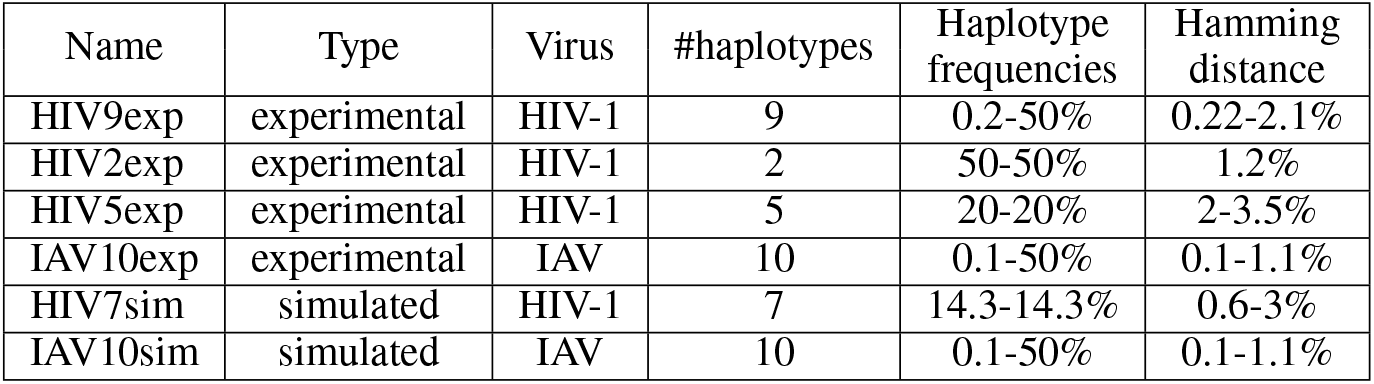
Four experimental and two simulated sequencing datasets of human immunodeficiency virus type 1 (HIV-1) and influenza A virus (IAV). The datasets contain MiSeq and PacBio reads from intra-host viral populations consisting of two to ten variants each with frequencies in the range of 0.1-50%, and Hamming distances between variants in the range of 0.1-3.5%.

| Name | Type | Virus | #haplotypes | Haplotype frequencies | Hamming distance |
| --- | --- | --- | --- | --- | --- |
| HIV9exp | experimental | HIV-1 | 9 | 0.2-50% | 0.22-2.1% |
| HIV2exp | experimental | HIV-1 | 2 | 50-50% | 1.2% |
| HIV5exp | experimental | HIV-1 | 5 | 20-20% | 2-3.5% |
| IAV10exp | experimental | IAV | 10 | 0.1-50% | 0.1-1.1% |
| HIV7sim | simulated | HIV-1 | 7 | 14.3-14.3% | 0.6-3% |
| IAV10sim | simulated | IAV | 10 | 0.1-50% | 0.1-1.1% |

To validate our method’s performance, we have introduced two novel in vitro sequencing HIV-1 benchmarks, which consist of Illumina MiSeq experiments on haplotype mixtures based on the mutation profile from an existing patient.

Finally, there is a essential gap in existing quality measures of intra-host viral population assembly. Up-to-date, instead of *populations* (i.e. haplotypes with their frequencies), only *sets* of reconstructed and the ground truth haplotypes are compared (29). Here we propose to measure differences between haplotype populations using Matching Error and the Earth Mover’s Distance which account for both the distances between haplotypes and their frequencies.

## MATERIALS AND METHODS

### CliqueSNV algorithm idea

A schematic diagram of the CliqueSNV algorithm is shown in Figure 1. The algorithm takes aligned reads as input and infers haplotype sequences with their frequencies as output. The method consists of six steps:

- Step 1 uses aligned reads to build the consensus sequence and identifies all SNVs. Then all pairs of SNVs are tested for dependency and are then divided into three groups: *linked, forbidden*, or *unclassified*. Each SNV is represented as a pair (*p,n*) of its position *p* and nucleotide value n in the aligned reads. If there are enough reads that have two SNVs (*p,n*) and (*p′,n′*) simultaneously, then they are tested for dependency. If the dependency test is positive and statistically significant (see CliqueSNV algorithm details for more information), then the algorithm classifies these two SNVs as *linked*. Otherwise, these two SNVs are tested for independency. If the independency test is positive and statistically significant (see Detailed description for details), then these two SNVs are classified as a *forbidden* pair.
- In Step 2, we build a graph *G* =(*V,E*) with a set of nodes V representing SNVs, and a set of edges E connecting linked SNV pairs.
- Ideally, SNVs of each true minority haplotype form a clique in G. A maximal clique *C*⊆*V* is a set of nodes such that (*u,v*) ∈ *E* for any *u,v* ∈ *C* and for any *x*∉*C* there is *u* ∈ *C* such that *(x,u)*∉*E* Step 3 finds all *maximal cliques* in *G*.
- For real sequencing data, the linkage between some SNV pairs may be undetected due to sequencing noise, uneven coverage, or the shortness of the NGS reads. As a result, a single clique corresponding to a haplotype will be split into several overlapping cliques. Step 4 merges such overlapping cliques. In order to avoid. if they contain a forbidden SNV pair.
- Step 5 assigns each read to a merged clique with which it shares the largest number of SNVs. Then CliqueSNV builds a consensus haplotype from all reads assigned to merging distinct haplotypes, two cliques are not merged a single merged clique.
- Finally, haplotype frequencies are estimated via an expectation-maximization algorithm in Step 6.

**Figure 1.**
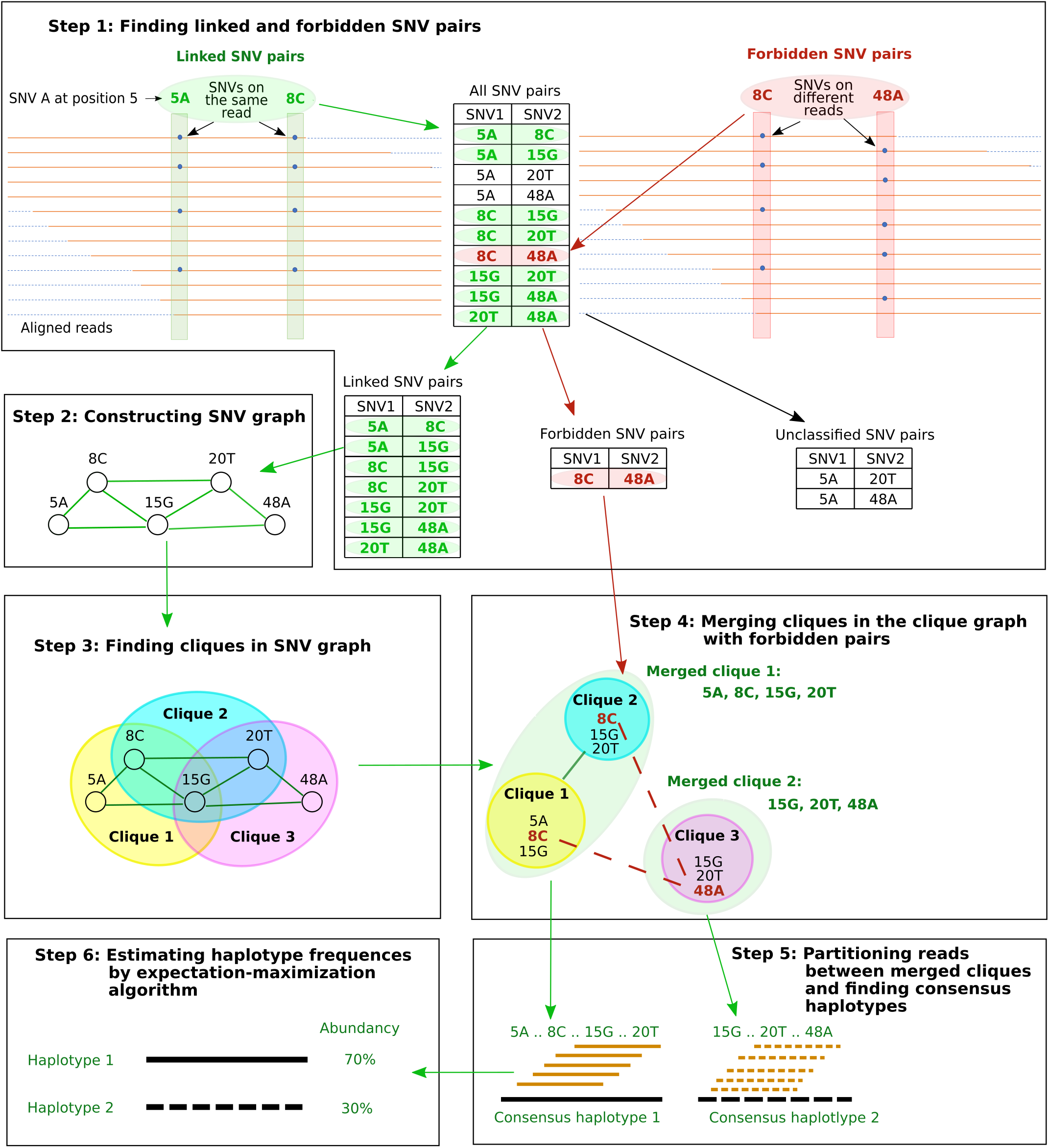
Schematic representation of the CliqueSNV algorithm. Where SNV is single nucleotide variation.

### Intra-host viral population sequencing benchmarks

We tested the ability of CliqueSNV to assemble haplotype sequences and estimate their frequencies from PacBio and MiSeq reads using four real (experimental) and two simulated datasets from HIV and IAV samples (Table 1). Each dataset contains between two to ten haplotypes with frequencies of 1 to 50%. The Hamming distances between pairs of variants for each dataset are shown in Figure S1.

#### Experimental datasets

(1-2). *HIV-1 subtype B plasmid mixtures and MiSeq reads (HIV2exp and HIV9exp)*. We designed nine in silico plasmid constructs comprising a 950-bp region of the HIV-1 subtype B polymerase (pol) gene that were then synthesized and cloned into pUCIDT-Amp (Integrated DNA Technologies, Skokie, IL). Each clone was confirmed by Sanger sequencing. This 950-bp region at the beginning of *pol* contains known protease and reverse transcriptase genes that are monitored for drug-resistant mutations and is monitored with sequence analysis for patient care. Each of these plasmids contains a specific set of point mutations chosen using mutation profiles of patient p7 from a real clinical study (53) to create nine unique synthetic HIV-1 *pol* haplotypes. Different proportions of these plasmids were mixed and then sequenced using an Illumina MiSeq protocol to obtain 2×300-bp reads (see Supplementary Methods). HIV2exp and HIV9exp are mixtures of two and nine variants, respectively.
3. *HIV-1 subtype B mixture and MiSeq reads (HIV5exp)*. This dataset consists of Illumina MiSeq 2 × 250-bp reads with an average read coverage of ^~^20,000 × obtained from a mixture of five HIV-1 isolates: 89.6, HXB2, JRCSF, NL43, and YU2 available at (52). Isolates have pairwise Hamming distances in the range from 2-3.5%(27 to 46-bp differences). The original HIV-1 sequence length was 9.3kb, but was reduced to the beginning of *pol* with a length of 1.3kb.
4. *Influenza A mixture and PacBio reads (IAV10exp)*. This benchmark contains ten IAV virus clones that were mixed at a frequency of 0.1-50%. The Hamming distances between clones ranged from 0.1-1.1% (2-22-bp differences) (48). The 2kb-amplicon was sequenced using the PacBio platform yielding a total of 33,558 reads with an average length of 1973 nucleotides.

#### Simulated datasets

1. *HIV-1 subtype B mixture and MiSeq reads (HIV7sim)*. This benchmark contains simulated Illumina MiSeq reads with a 10k-coverage of 1-kb *pol* sequences. The reads were simulated from seven equally distributed HIV-1 variants chosen from the NCBI database: AY835778, AY835770, AY835771, AY835777, AY835763, AY835762, and AY835757. The Hamming distances between clones are in the range from 0.6-3.0% (6 to 30-bp differences). We used SimSeq (54) for generating reads.
2. *Influenza A mixture and MiSeq reads (IAV10sim)*. This benchmark contains simulated IAV Illumina MiSeq reads with the same IAV haplotypes and their frequencies as for the IAV10exp benchmark. The sequencing of a 2kb-amplicon with 40k coverage with paired Illumina MiSeq reads was simulated by SimSeq (54) with the default sequencing error profile in SimSeq.

### Validation metrics for viral population inference

#### Precision and recall

Inference quality is typically measured by precision and recall.

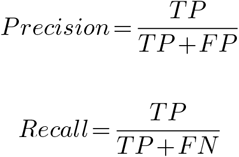

where *TP* is the number of true predicted haplotypes, *FP* is the number of false predicted haplotypes, and *FN* is the number of undiscovered haplotypes.

Initially we measured precision and recall strictly by treating a predicted haplotype with a single mismatch as an *FP*. Additionally, like in (29) we introduced an acceptance threshold, which is the number of mismatches permitted for a predicted haplotype to count as a TP.

#### Matching errors between populations

However, precision and recall do not take into account (i) distances between true and inferred viral variants as well as (ii) the frequencies of the true and inferred viral variants. Instead, we chose to use analogues of precision and recall defined for populations as follows.

Let *T*={(*t,f_t_*)}, be the true haplotype population, where *f_t_* is the frequency of the true haplotype *t*, Σ_*t*∈*T*_ *f_t_* = 1.

Similarly, let *P* = {(*p,f_p_*)}, be the reconstructed haplotype population, where *f_p_* is the frequency of the reconstructed haplotype *p*, Σ_*p*∈*P*_ *f_p_* = 1. Let *d_pt_* be the distance between haplotypes p and t. Thus, instead of precision, we used the *matching error E_T→P_* which measures how well each reconstructed haplotype *p*∈*P* weighted by its frequency is matched by the closest true haplotype.

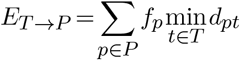

Indeed, precision increases while *E_T→P_* decreases and reaches 100% when *E_T→P_* = 0. Similarly, instead of recall, we propose to use the *matching error E_T←P_* which measures how well each true haplotype *t*∈*T* weighted by its frequency is matched by the closest reconstructed haplotype. (55)

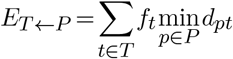

Note that recall increases while *E_T←P_* decreases and reaches 100% when *E_T←P_* = 0.

#### Earth mover’s distance (EMD) between populations

The matching errors described above match haplotypes of true and reconstructed populations but do not match their frequencies. In order to simultaneously match haplotype sequences and their frequencies, we allowed for a fractional matching when portions of a single haplotype *p* of population *P* are matched to portions of possibly several haplotypes of *T* and *vice versa.* Thus, we separated *f_p_* into *f_pt_’s* each denoting portion of *p* matched to *t* such that *f_p_* = Σ_*t*∈*T*_ *f_pt_*, *f_pt_* ≥ 0. Symmetrically, *f_t_’s* are also separated into *f_pt_’s,* i.e, Σ_*p*∈*P*_ *f_pt_* = *f_t_*. Finally, we chose *f_pt_’s* minimizing the total error of matching *T* to *P* which is also known as Wasserstein metric or the EMD between *T* and *P* (56, 57).

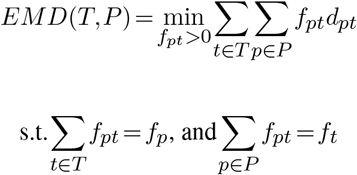

EMD is efficiently computed as an instance of the transportation problem using network flows.

EMDs can vary a lot over different benchmarks since they may have different complexities, which depends on the number of true variants, the frequency distribution, the similarity between haplotypes, sequencing depth, sequencing error rate, and many other parameters. Hence, we measured the complexity of a benchmark as the EMD between the true population and a population consisting of a single consensus haplotype (58).

### CliqueSNV algorithm details

Data input for CliqueSNV consists of PacBio or Illumina reads from an intra-host viral population aligned to a reference genome. Output is the set of inferred viral variant RNA sequences with their frequencies. The formal high-level pseudocode of the CliqueSNV algorithm is described in the supplementary materials. Below we describe in detail the six major steps of CliqueSNV that are schematically presented in Figure 1.

#### Step 1: Finding linked and forbidden SNV pairs

At a given genomic position *I*, the most frequent nucleotide is referred to as a *major variant* and is denoted 1. Let us fix one of the less frequent nucleotide (referred to as a *minor variant*) and denote it 2. A pair of variants at two distinct genomic positions *I* and *J* is referred to as a 2-haplotype. There are four 2-haplotypes with major and minor variants at *I* and *J*: (11),(12),(21), and (22). Let *O*_11_, *O*_12_, *O*_21_, *O*_22_ be the observed counts of 2-haplotypes in the reads covering *I* and *J*. In this step, CliqueSNV tries to decide whether the *O*_22_ reads are sequencing errors or they are produced by an existing haplotype containing the 2-haplotype (22).

The pairs of minor variants (referred to as SNV pairs) are classified into three categories: linked, forbidden, and unclassified. An SNV pair is *linked* if it is highly probable that there exists a haplotype containing both minor variants. On the contrary, an SNV *pair* is *forbidden* if it is extremely unlikely that the corresponding minor variants belong to the same haplotype. All other SNV pairs are referred to as *unclassified*.

Assuming that errors are random, it has been proven in (59) that if the 2-haplotype (22) does not exist, then the expected number of reads *E*_22_ containing the 2-haplotype (22) should not exceed

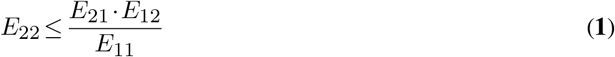

where *E*_21_, *E*_12_, and *E*_11_ are the expected numbers of reads containing the 2-haplotypes (21), (12) and (11), respectively. To determine if a pair of SNVs (the minor variants in positions *I* and *J*) are linked, we need to estimate the probability that the observed counts of 2-haplotypes *O*_11_, *O*_12_, *O*_21_, *O*_22_ are produced by 2-haplotype counts satisfying equation **1**.

Let *n* be the total number of reads covering both positions *I* and *J*. Then

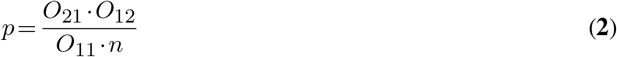

is the probability of observing *O*_22_ reads with the both minor variants given that the variant (22) does not exist.

The 2-haplotype (22) exists with high probability 1 − *P* and the corresponding pair of SNVs is linked if the value of *p* satisfies the following inequality (59)

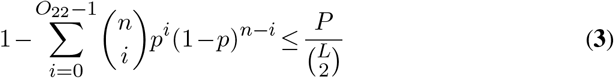

where *P* is the user-defined *P*-value (by default *P* = 0.01) and dividing by 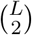 is the Bonferroni correction for multiple testing.

Pairs of SNVs passing this linkage test are classified as a linked SNV pairs. For every other pair of SNVs, we check whether they can be classified as a *forbidden* SNV pair, i.e., whether the probability of observing at most 0_22_ reads is low enough (<0.05) given that the variant (22) has frequency *T*_22_≥*t* (by default *t*=0.001).

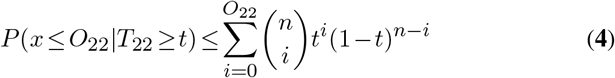

#### Step 2: Constructing the SNV graph

The SNV graph *G* = (*V,E*) consists of vertices corresponding to minor variants and edges corresponding to linked pairs of minor variants from different positions. If the intra-host population consists of very similar haplotypes, then graph *G* is very sparse. Indeed, the PacBio dataset for IAV encompassing 2,500 positions is split into 10,000 vertices, while the SNV graph contains only 700 edges, and, similarly, the simulated Illumina read dataset for the same haplotypes contains only 368 edges.

Note that the isolated minor variants correspond to genotyping errors unless they have a significant frequency. This fact allows us to estimate the number of errors per read, assuming that all isolated SNVs are errors. As expected, the distribution of the PacBio reads has a heavy tail (see Figure S4), which implies that most reads are (almost) error free, while a small number of heavy-tail reads accumulate most of the errors. Our analysis allows the identification of such reads, which can then be filtered out. By default, we filter out ≈10% of PacBio reads, but we do not filter out any Illumina reads. The SNV graph is then constructed for the reduced set of reads. Such filtering allows the reduction of systematic errors and refines the SNV graph significantly.

#### Step 3: Finding cliques in the SNV graph *G*

Although the MAX CLIQUE is a well-known NP-complete problem and there may be an exponential number of maximal cliques in *G*, a standard Bron-Kerbosch algorithm requires little computational time since *G* is very sparse (60).

#### Step 4: Merging cliques in the clique graph *C_G_*

The clique graph *C_G_*= (*C,F,L*) consists of vertices corresponding to cliques in the SNV graph *G* and two sets of edges *F* and *L. A* forbidding edge (*p,q*)∈*F* connects two cliques *p* and *q* with at least one forbidden pair of minor variants from *p* and *q* respectively. A *linking edge* (*p,q*)∈*L* connects two cliques *p* and *q*, (*p,q*)∉*F*, with at least one linked pair of minor variants from *p* and *q* respectively. Any true haplotype corresponds to a maximal (*L\F*)-connected subgraph *H* of *C_G_* which is connected with edges from *L* and does not contain any edge from *F* (see Fig. 1 (4)).

Unfortunately, even deciding whether there is a *L*-path between *p* and *q* avoiding forbidding edges is known to be NP-hard (61). We find all subgraphs *H* as follows (see Figure S5): (i) connect all pairs of vertices except connected with forbidding edges, (ii) find all maximal super-cliques in the resulted graph 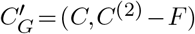 using (60), (iii) split each super-clique into *L*-connected components, and (iv) filter out the *L*-connected components which are proper subsets of other maximal *L*-connected components.

#### Step 5: Partitioning reads between merged cliques and finding consensus haplotypes

Let *S* be the set of all positions containing at least one minor variant in *V*. Let qs be an *major clique* corresponding to a haplotype with all major variants in *S*. The distance between a read *r* and a clique q equals the number of variants in *q* that are different from the corresponding nucleotides in *r*. Each read *r* is assigned to the closest clique *q* (which can possibly be *q_S_*). In case of a tie, we assign *r* to all closest cliques.

Finally, for each clique *q*, CliqueSNV finds the consensus *v(q*) of all reads assigned to *q*. Then *v(q*) is extended from *S* to a full-length haplotype by setting all non-*S* positions to major SNVs.

#### Step 6: Estimating haplotype frequencies by using the expectation-maximization (EM) algorithm

CliqueSNV estimates the frequencies of the assembled intra-host haplotypes via an expectation-maximization algorithm similar to the one used in IsoEM (62). Let *K* be the number of assembled viral variants, and let *α* be the probability of sequencing error. EM algorithm works as follows:

1. Initialize frequencies of viral variants 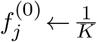, Compute the probability of *l_i_*-long read *r_i_* = 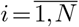, being emitted by viral variant, 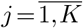, 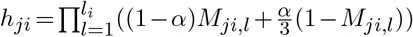 where *M_ji,l_*-indicator if *i*-th read coincides with *j*-th viral variant in the position *l*
2. (Expectation) Update the amount of read *r_i_* emitted by the *j*th viral variant 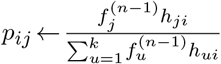
3. (Maximization) Update the frequency of the *j*th viral variant 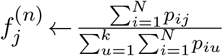
4. if 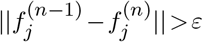, then *n←n*+1 and go to step 2
5. Output estimated frequencies *f^(n)^*

## RESULTS

### Performance of haplotyping methods

We compared CliqueSNV to the 2SNV, PredictHaplo, and aBayesQR haplotyping methods. Since CliqueSNV, PredictHaplo and aBayesQR use Illumina reads, we compared them using the HIV9exp, HIV2exp, HIV5exp, HIV7sim, and IAV10sim datasets. Since CliqueSNV, 2SNV, and PredictHaplo can also use PacBio reads, we compared them using the IAV10exp dataset. We also used consensus sequences in the comparisons (58) because of its simplicity and to evaluate sequences most similar to those generated by the Sanger sequencing method (63).

The precision and recall of haplotype discovery for each method is provided in Table 2. CliqueSNV had the best precision and recall for five of the six datasets. For the HIV5exp dataset, PredictHaplo was more conservative and predicted less false positive variants (better precision) than CliqueSNV.

**Table 2.** Prediction statistics of haplotype reconstruction methods using experimental and simulated (a) MiSeq and (b) PacBio datasets. The precision and recall was evaluated stringently such that if a predicted haplotype has at least one mismatch to its closest answer, then that haplotype is scored as a false positive.

| Benchmark | CliqueSNV |  | aBayesQR |  | PredictHaplo |  |
| --- | --- | --- | --- | --- | --- | --- |
|  | Precision | Recall | Precision | Recall | Precision | Recall |
| HIV9exp | **0.60** | **0.33** | 0.00 | 0.00 | 0.00 | 0.00 |
| HIV2exp | **0.66** | **1.00** | 0.11 | 0.50 | 0.50 | 0.50 |
| HIV5exp | 0.18 | **0.40** | 0.00 | 0.00 | **0.33** | 0.20 |
| HIV7sim | **1.00** | **0.71** | **1.00** | 0.42 | 0.45 | **0.71** |
| IAV10sim | **0.75** | **0.30** | 0.11 | 0.10 | 0.33 | 0.10 |

(a)
| Benchmark | CliqueSNV |  | 2SNV |  | PredictHaplo |  |
| --- | --- | --- | --- | --- | --- | --- |
|  | Precision | Recall | Precision | Recall | Precision | Recall |
| IAV10exp | **1.00** | **1.00** | 0.82 | 0.90 | 0.70 | 0.70 |

Following study (29), we also showed how precision and recall grew with the reduction of restriction on mismatches (Fig. 2). The number of true predicted haplotypes for CliqueSNV was always greater than that of the other methods on real experimental sequencing benchmarks indicating that CliqueSNV more accurately identified the true haplotypes. The number of falsely predicted haplotypes for CliqueSNV was always lower than those for aBayesQR, but similar to those predicted by PredictHaplo on four out of five datasets indicating that both CliqueSNV and PredictHaplo had the best precision with MiSeq datasets.

**Figure 2.**
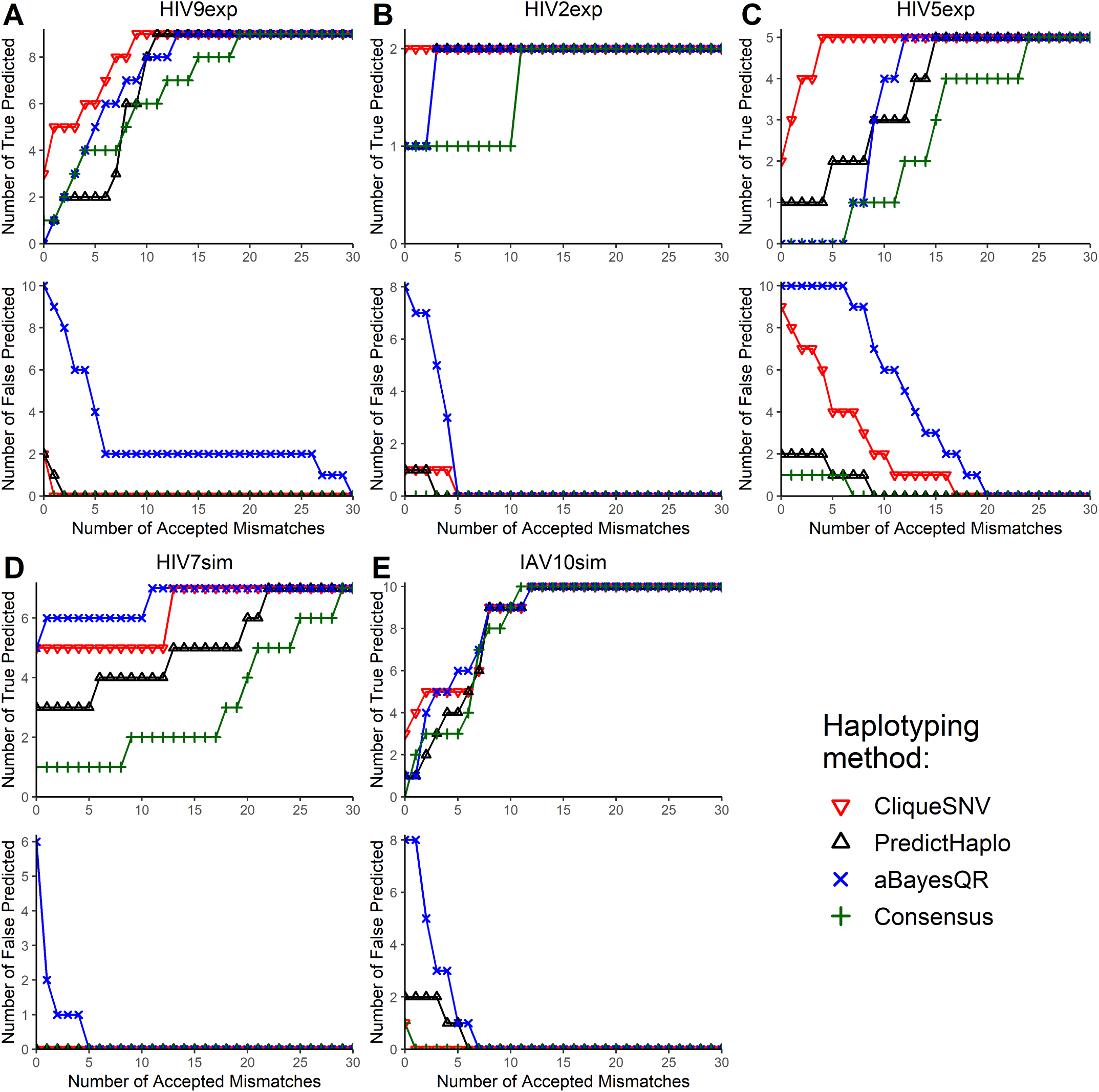
The number of true and false predicted haplotypes depending on the number of accepted mismatches for five benchmarks: (A) HIV9exp; (B) HIV2exp; (C) HIV5exp; (D) HIV7sim; (E) IAV10sim. Two haplotypes are regarded identical if the Hamming distance between them is at most the number of accepted mismatches.

Matching distance analysis showed that matching distances *E_T←P_* and *E_T→P_* are better for CliqueSNV than for both PredictHaplo and aBayesQR on four out of five MiSeq datasets (Fig. 3). For HIV7sim, *E_T←P_* for aBayesQR was slightly better than for CliqueSNV. Using HIV9exp, HIV2exp, HIV7sim, and IAV10sim datasets, the *E_T←P_* and *E_T→P_* for CliqueSNV were very close to zero indicating that the predictions were almost perfect. Since *E_T←P_* and *E_T→P_* correlate with precision and recall, matching distance analysis indicates that CliqueSNV had a better precision, and significantly outperformed both PredictHaplo and aBayesQR. Since aBayesQR had a higher *E_T→P_* on MiSeq datasets, it is more likely to make more false predictions. Notably, on the HIV7sim dataset, aBayesQR outperformed both CliqueSNV and PredictHaplo by *E_T←P_*.

**Figure 3.**
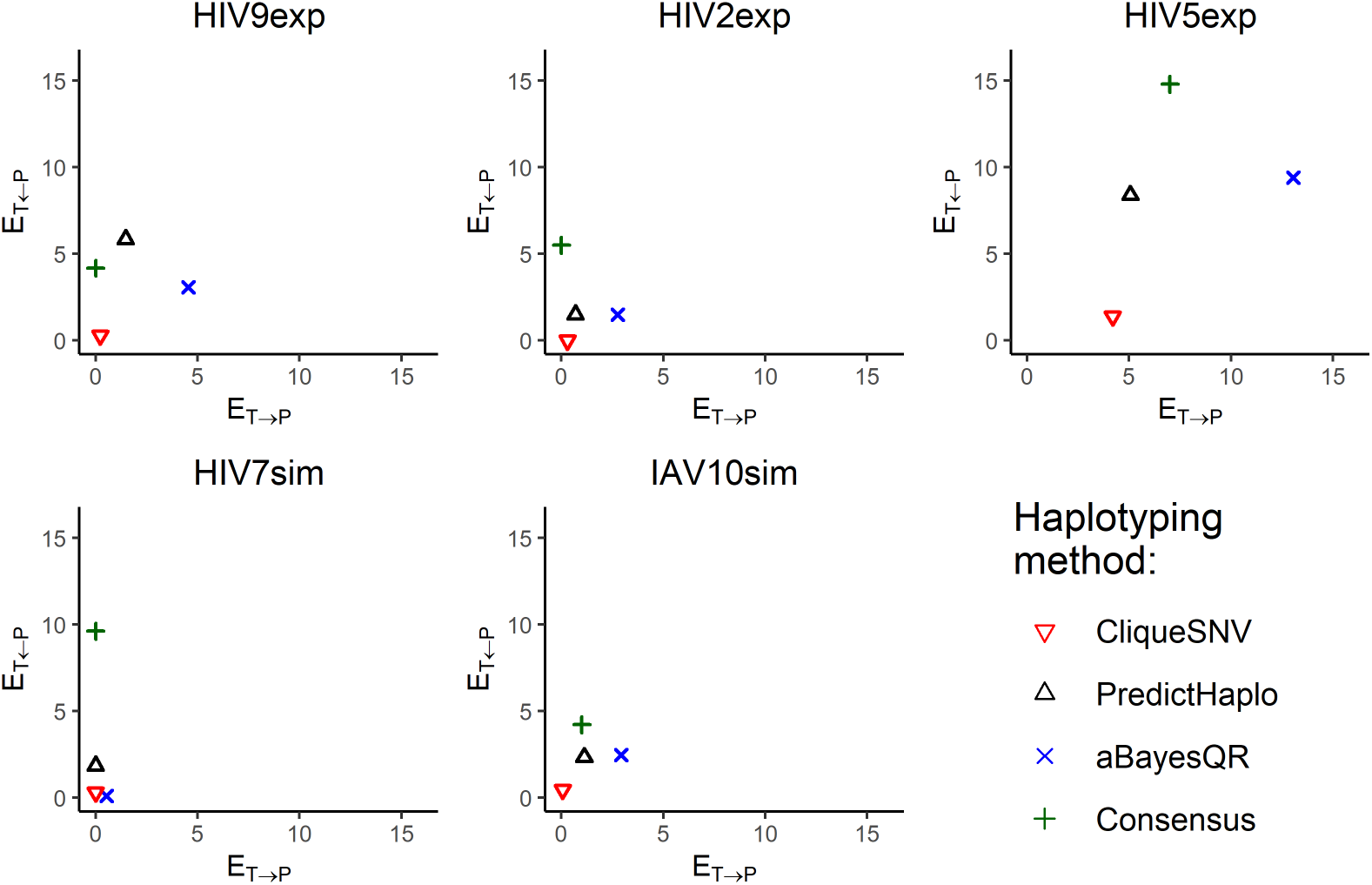
Matching distances *E_T←P_* and *E_T→P_* between the true haplotype population *T* and the reconstructed haplotype population *P* for five benchmarks.

The EMD between the predicted and true haplotype populations for all five MiSeq datasets are shown in Figure 4. The exact EMD values are provided in Table 3. CliqueSNV provided the lowest (the best) EMD across all tools on four out of five MiSeq benchmarks. For the simulated and PacBio datasets, CliqueSNV had almost a zero EMD indicating a low error in predictions. PredictHaplo had a lower EMD than aBayesQR on four out of five MiSeq datasets. aBayesQR has almost a zero EMD with the HIV7sim dataset and outperformed CliqueSNV, while using the HIV5exp dataset, aBayesQR performed poorer than other methods.

**Figure 4.**
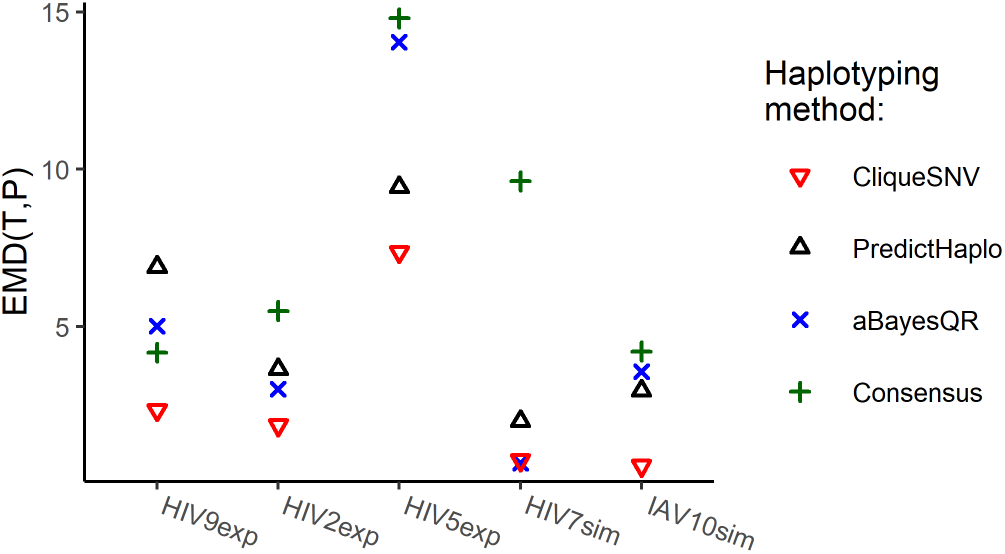
Earth Movers’ Distance (EMD) between true and reconstructed haplotype populations for five benchmarks.

**Table 3.** Earth Movers’ Distance from predicted haplotypes to the true haplotype population and haplotyping method improvement. Four haplotyping methods(aBayesQR, CliqueSNV, Consensus, PredictHaplo) are benchmarked using five MiSeq (a) and one PacBio datasets (b). The column Impr. (improvement) shows how much better is prediction of haplotyping method over inferred consensus, and it is calculated as 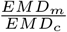, where *EMD_c_* is an EMD for consensus, and *EMD_m_* is an EMD for method.

| Benchmark | Consensus | CliqueSNV |  | aBayesQR |  | PredictHaplo |  |
| --- | --- | --- | --- | --- | --- | --- | --- |
|  | EMD | EMD | Impr. | EMD | Impr. | EMD | Impr. |
| HIV9exp | 4.18 | **2.35** | **1.78** | 5.02 | 0.83 | 6.90 | 0.61 |
| HIV2exp | 5.50 | **1.87** | **2.94** | 3.02 | 1.82 | 3.65 | 1.51 |
| HIV5exp | 14.80 | **7.37** | **2.01** | 14.05 | 1.05 | 9.43 | 1.57 |
| HIV7sim | 9.63 | 0.76 | 12.72 | **0.67** | **14.4** | 2.00 | 4.80 |
| IAV10sim | 4.22 | **0.59** | **7.2** | 3.57 | 1.18 | 2.97 | 1.42 |

(a)
| Benchmark | Consensus | CliqueSNV |  | 2SNV |  | PredictHaplo |  |
| --- | --- | --- | --- | --- | --- | --- | --- |
|  | EMD | EMD | Impr. | EMD | Impr. | EMD | Impr. |
| IAV10exp | 4.22 | **0.22** | **19.18** | 0.23 | 18.35 | 0.38 | 11.12 |

Next, CliqueSNV, 2SNV, and PredictHaplo were compared using the IAV10exp benchmark dataset (see Table S1). CliqueSNV correctly recovered all ten true variants, including the haplotype with frequencies significantly below the sequencing error rate. 2SNV recovered nine true variants but found one false positive. PredictHaplo recovered only seven true variants and falsely predicted three variants. To further explore the precision of these three methods with the IAV10exp data, we simulated low-coverage datasets by randomly subsampling *n* = 16*K*,8*K*,4*K* reads from the original data. for each dataset, CliqueSNV found at least one true variant more than both 2snv and predicthaplo.

### Runtime comparison

To compare the computational run time of each method, we used the same PC (Intel(R) Xeon(R) CPU X5550 2.67GHz x2 8 cores per CPU, DIMM DDR3 1,333 MHz RAM 4Gb x12) with the CentOS 6.4 operating system. The runtime of CliqueSNV is sublinear with respect to the number of reads while the runtime of PredictHaplo and 2SNV exhibit super-linear growth. For the 33k IAV10sim reads the CliqueSNV analysis took 21 seconds, while PredictHaplo and 2SNV took around 30 minutes. The runtime of CliqueSNV is quadratic with respect to the number of SNVs rather than by the length of the sequencing region (Fig. S2).

We also generated five HIV-1 variants within 1% Hamming distance from each other, which is the estimated genetic distance between related HIV variants from the same person (64). Then we simulated 1M Illumina reads for sequence regions of length 566, 1132, 2263 and 9181 nucleotides for which CliqueSNV required 37, 144, 227, and 614 seconds, respectively, for analyzing these datasets (Fig. S3). For the HIV2exp benchmark, aBayesQR, PredictHaplo, and CliqueSNV required over ten hours, 24 minutes, and only 79 seconds, respectively.

## DISCUSSION

Assembly of haplotype populations from noisy NGS data is one of the most challenging problems of computational genomics. High-throughput sequencing technologies, such as Illumina MiSeq and HiSeq, provide deep sequence coverage that allows discovery of rare, clinically relevant haplotypes. However, the short reads generated by the Illumina technology require assembly that is complicated by sequencing errors, an unknown number of haplotypes in a sample, and the genetic similarity of haplotypes within a sample. Furthermore, the frequency of sequencing errors in

Illumina reads is comparable to the frequencies of true minor mutations (41). The recent development of single-molecule sequencing platforms such as PacBio produce reads that are sufficiently long to span entire genes or small viral genomes. Nonetheless, the error rate of single-molecule sequencing is exceptionally high reaching 13 —14% (65), which hampers PacBio sequencing to detect and assemble rare viral variants.

We developed CliqueSNV, a new reference-based assembly method for reconstruction of rare genetically-related viral variants such as those observed during infection with rapidly evolving RNA viruses like HIV, HCV and IAV. We demonstrated that CliqueSNV infers accurate haplotyping in the presence of high sequencing error rates and is also suitable for both single-molecule and short-read sequencing. In contrast to other haplotyping methods, CliqueSNV infers viral haplotypes by detection of clusters of statistically linked SNVs rather than through assembly of overlapping reads used with methods such as Savage (24).

Applied to the novel in vitro sequencing HIV-1 benchmark, CliqueSNV correctly reconstructed 87% of the intra-host haplotype population. At the same time, other state-of-the-art tools were not able to recover even a single haplotype without errors. Additionally, we have used the only previously known and commonly used in vitro benchmark (26) and simulated datasets to evaluate the accuracy of existing haplotyping methods. In contrast to the existing methods, CliqueSNV was able to detect minority haplotypes at a low 0.1% frequency and distinguish minority haplotypes differently in only two base pairs.

Although very accurate and fast, CliqueSNV has some limitations. Unlike Savage (24), CliqueSNV is not a *de novo* assembly tool and requires a reference viral genome. This obstacle could easily be addressed by using Vicuna (58) or other analogous tools to first assemble a consensus sequence from the NGS reads, which can then be used as a reference. Another limitation is for variants that differ only by isolated SNVs separated by long conserved genomic regions longer than the read length which may not be accurately inferred by CliqueSNV. While such situations usually do not occur for viruses, where mutations are typically densely concentrated in different genomic regions, we plan to address this limitation in the next version of CliqueSNV.

The ability to accurately infer the structure of intra-host viral populations makes CliqueSNV applicable for studying viral evolution, transmission and examining the genomic compositions of RNA viruses. In addition, we envision that the application of our method could be extended to other highly heterogeneous genomic populations, such as metagenomes, immune repertoires, and cancer cell genes.

## DATA AVAILABILITY

The datasets HIV2exp and HIV9exp have been deposited in the Sequence Read Archive under accession number SRR12042289 and SRR12042290, respectively.

The links to the data sets and the consensus sequences of the individual strains are available at https://github.com/Sergey-Knyazev/CliqueSNV-validation/blob/master/relevant_haplotypes/HIV9exp.fasta

## Software availability

CliqueSNV is available at https://github.com/vtsyvina/CliqueSNV

## Validation scripts availability

All scripts and configuration files that were used for validation of the tools are available at https://github.com/Sergey-Knyazev/CliqueSNV-validation

## ACKNOWLEDGEMENTS

Use of trade names is for identification only and does not imply endorsement by the U.S. Department of Health and Human Services, the Public Health Service, or the Centers for Disease Control and Prevention (CDC). The findings and conclusions in this paper are those of the authors and do not necessarily represent the official position of the CDC.

## REFERENCES

1. Kilmarx, P. H. (2009) Global epidemiology of HIV. Current Opinion in HIV and AIDS, 4(4), 240–246.

2. Hajarizadeh, B., Grebely, J., and Dore, G. J. (2013) Epidemiology and natural history of HCV infection. Nature Reviews Gastroenterology and Hepatology, 10(9), 553–562.

3. Lozano, R., Naghavi, M., Foreman, K., Lim, S., Shibuya, K., Aboyans, V., Abraham, J., Adair, T., Aggarwal, R., Ahn, S. Y., et al. (2012) Global and regional mortality from 235 causes of death for 20 age groups in 1990 and 2010: a systematic analysis for the Global Burden of Disease Study 2010. The lancet, 380(9859), 2095–2128.

4. Eigen, M., McCaskill, J., and Schuster, P. (1989) The molecular quasi-species. Advances in chemical physics, 75, 149–263.

5. Martell, M., Esteban, J., Quer, J., Genesca, J., Weiner, A., Esteban, R., Guardia, J., and Gomez, J. (1992) Hepatitis C virus (HCV) circulates as a population of different but closely related genomes: quasispecies nature of HCV genome distribution. Journal of Virology, 66 pp. 3225–3229.

6. Steinhauer, D. and Holland, J. (1987) Rapid evolution of RNA viruses. Annual Review of Microbiology, 41 pp. 409–433.

7. Domingo, E., Sheldon, J., and Perales, C. (2012) Viral quasispecies evolution. Microbiology and Molecular Biology Reviews, 76(2), 159–216.

8. Rodriguez-Frias, F., Buti, M., Tabernero, D., and Homs, M. (2013) Quasispecies structure, cornerstone of hepatitis B virus infection: mass sequencing approach. World J Gastroenterol, 19(41), 6995–7023.

9. Xu, D., Zhang, Z., and Wang, F.-S. (March, 2004) SARS-Associated Coronavirus Quasispecies in Individual Patients. New England Journal of Medicine, 350(13), 1366–1367.

10. Shen, Z., Xiao, Y., Kang, L., Ma, W., Shi, L., Zhang, L., Zhou, Z., Yang, J., Zhong, J., Yang, D., Guo, L., Zhang, G., Li, H., Xu, Y., Chen, M., Gao, Z., Wang, J., Ren, L., and Li, M. (March, 2020) Genomic Diversity of Severe Acute Respiratory Syndrome–Coronavirus 2 in Patients With Coronavirus Disease 2019. Clinical Infectious Diseases,.

11. Beerenwinkel, N., Sing, T., Lengauer, T., Rahnenfuehrer, J., and Roomp, K. (2005) Computational methods for the design of effective therapies against drug resistant HIV strains. Bioinformatics, 21, 3943–3950.

12. Douek DC, Kwong PD, N. G. (2006) The rational design of an AIDS vaccine. Cell, 124, 677–681.

13. Gaschen, B., Taylor, J., Yusim, K., Foley, B., and Gao, F. (2002) Diversity considerations in HIV-1 vaccine selection. Science, 296, 2354–2360.

14. Holland, J., De La Torre, J., and Steinhauer, D. (1992) RNA virus populations as quasispecies. Curr Top Microbiol Immunol, 176, 1–20.

15. Rhee, S.-Y., Liu, T., Holmes, S., and Shafer, R. (2007) HIV-1 Subtype B Protease and Reverse Transcriptase Amino Acid Covariation. PLoS Comput Biol, 3, e87.

16. Campo, D. S., Skums, P., Dimitrova, Z., Vaughan, G., Forbi, J. C., Teo, C.-G., Khudyakov, Y., and Lau, D. T. (2014) Drug Resistance of a Viral Population and Its Individual Intrahost Variants During the First 48 Hours of Therapy. Clinical Pharmacology & Therapeutics, 95(6), 627–635.

17. Skums, P., Bunimovich, L., and Khudyakov, Y. (2015) Antigenic cooperation among intrahost HCV variants organized into a complex network of cross-immunoreactivity. Proceedings of the National Academy of Sciences, 112(21), 6653–6658.

18. Campo, D. S., Xia, G.-L., Dimitrova, Z., Lin, Y., Forbi, J. C., Ganova-Raeva, L., Punkova, L., Ramachandran, S., Thai, H., Skums, P., et al. (2016) Accurate genetic detection of hepatitis C virus transmissions in outbreak settings. Journal of Infectious Diseases, 213(6), 957–965.

19. Glebova, O., Knyazev, S., Melnyk, A., Artyomenko, A., Khudyakov, Y., Zelikovsky, A., and Skums, P. (December, 2017) Inference of genetic relatedness between viral quasispecies from sequencing data. BMC Genomics, 18(S10).

20. Skums, P., Zelikovsky, A., Singh, R., Gussler, W., Dimitrova, Z., Knyazev, S., Mandric, I., Ramachandran, S., Campo, D., Jha, D., Bunimovich, L., Costenbader, E., Sexton, C., O’Connor, S., Xia, G.-L., and Khudyakov, Y. (June, 2017) QUENTIN: reconstruction of disease transmissions from viral quasispecies genomic data. Bioinformatics, 34(1), 163–170.

21. Wymant, C., Hall, M., Ratmann, O., Bonsall, D., Golubchik, T., de Cesare, M., Gall, A., Cornelissen, M., and and, C. F. (November, 2017) PHYLOSCANNER: Inferring Transmission from Within- and Between-Host Pathogen Genetic Diversity. Molecular Biology and Evolution, 35(3), 719–733.

22. Melnyk, A., Knyazev, S., Vannberg, F., Bunimovich, L., Skums, P., and Zelikovsky, A. (May, 2019) Using Earth Mover’s Distance for Viral Outbreak Investigations. bioRxiv,.

23. Boskova, V. and Stadler, T. (June, 2020) PIQMEE: Bayesian phylodynamic method for analysis of large datasets with duplicate sequences. Molecular Biology and Evolution,.

24. Baaijens, J. A., El Aabidine, A. Z., Rivals, E., and Schönhuth, A. (2017) De novo assembly of viral quasispecies using overlap graphs. Genome Research, 27(5), 835–848.

25. Döring, M., Büch, J., Friedrich, G., Pironti, A., Kalaghatgi, P., Knops, E., Heger, E., Obermeier, M., Däumer, M., Thielen, A., Kaiser, R., Lengauer, T., and Pfeifer, N. (May, 2018) geno2pheno[ngs-freq]: a genotypic interpretation system for identifying viral drug resistance using next-generation sequencing data. Nucleic Acids Research, 46(W1), W271–W277.

26. Giallonardo, F. D., Töpfer, A., Rey, M., Prabhakaran, S., Duport, Y., Leemann, C., Schmutz, S., Campbell, N. K., Joos, B., Lecca, M. R., Patrignani, A., Däumer, M., Beisel, C., Rusert, P., Trkola, A., Günthard, H. F., Roth, V., Beerenwinkel, N., and Metzner, K. J. (June, 2014) Full-length haplotype reconstruction to infer the structure of heterogeneous virus populations. Nucleic Acids Research, 42(14), e115–e115.

27. Knyazev, S., Hughes, L., Skums, P., and Zelikovsky, A. (June, 2020) Epidemiological data analysis of viral quasispecies in the next-generation sequencing era. Briefings in Bioinformatics,.

28. Office of the Commissioner FDA authorizes marketing of first next-generation sequencing test for detecting HIV-1 drug resistance mutations. https://www.fda.gov/news-events/press-announcements/fda-authorizes-marketing-first-next-generation-sequencing-test-detecting-hiv-1-drug-resistance (May, 2019) Accessed: 2019-12-28.

29. Prabhakaran, S., Rey, M., Zagordi, O., Beerenwinkel, N., and Roth, V. (2014) HIV haplotype inference using a propagating Dirichlet process mixture model. IEEE/ACM Transactions on Computational Biology and Bioinformatics (TCBB), 11(1), 182–191.

30. Ahn, S. and Vikalo, H. (2017) aBayesQR: A Bayesian method for reconstruction of viral populations characterized by low diversity. In International Conference on Research in Computational Molecular Biology Springer pp. 353–369.

31. Töpfer, A., Zagordi, O., Prabhakaran, S., Roth, V., Halperin, E., and Beerenwinkel, N. (2013) Probabilistic inference of viral quasispecies subject to recombination. Journal of Computational Biology, 20(2), 113–123.

32. Töpfer, A., Marschall, T., Bull, R. A., Luciani, F., Schönhuth, A., and Beerenwinkel, N. (2014) Viral Quasispecies Assembly via Maximal Clique Enumeration. PLoS Computational Biology, 10(3).

33. Mangul, S., Wu, N. C., Mancuso, N., Zelikovsky, A., Sun, R., and Eskin, E. (2014) Accurate viral population assembly from ultra-deep sequencing data. Bioinformatics, 30(12), i329–i337.

34. Skums, P., Mancuso, N., Artyomenko, A., Tork, B., Mandoiu, I., Khudyakov, Y., and Zelikovsky, A. (2013) Reconstruction of viral population structure from next-generation sequencing data using multicommodity flows. BMC bioinformatics, 14(Suppl 9), S2.

35. Mancuso, N., Tork, B., Skums, P., Ganova-Raeva, L., Măndoiu, I., and Zelikovsky, A. (2011) Reconstructing viral quasispecies from NGS amplicon reads. In silico biology, 11(5), 237–249.

36. Zagordi, O., Bhattacharya, A., Eriksson, N., and Beerenwinkel, N. (2011) ShoRAH: estimating the genetic diversity of a mixed sample from next-generation sequencing data. BMC bioinformatics, 12(1), 119.

37. Astrovskaya, I., Tork, B., Mangul, S., Westbrooks, K., Măndoiu, I., Balfe, P., and Zelikovsky, A. (2011) Inferring viral quasispecies spectra from 454 pyrosequencing reads. BMC bioinformatics, 12(Suppl 6), S1.

38. Prosperi, M. C. and Salemi, M. (2012) QuRe: software for viral quasispecies reconstruction from next-generation sequencing data. Bioinformatics, 28(1), 132–133.

39. Zagordi, O., Töpfer, A., Prabhakaran, S., Roth, V., Halperin, E., and Beerenwinkel, N. (2012) Probabilistic inference of viral quasispecies subject to recombination. In Proceedings of the 16th Annual international conference on Research in Computational Molecular Biology Berlin, Heidelberg: Springer-Verlag RECOMB’12 pp. 342–354.

40. Skums, P., Dimitrova, Z., Campo, D. S., Vaughan, G., Rossi, L., Forbi, J. C., Yokosawa, J., Zelikovsky, A., and Khudyakov, Y. (2012) Efficient error correction for next-generation sequencing of viral amplicons. BMC Bioinformatics, 13(S-10), S6.

41. Skums, P., Artyomenko, A., Glebova, O., Campo, D. S., Dimitrova, Z., Zelikovsky, A., and Khudyakov, Y. (2016) ERROR CORRECTION OF NGS READS FROM VIRAL POPULATIONS. Computational Methods for Next Generation Sequencing Data Analysis,.

42. Barik, S., Das, S., and Vikalo, H. (2016) Viral Quasispecies Reconstruction via Correlation Clustering. bioRxiv, p. 096768.

43. Westbrooks, K., Astrovskaya, I., Campo, D., Khudyakov, Y., Berman, P., and Zelikovsky, A. (2008) HCV quasispecies assembly using network flows. Bioinformatics Research and Applications, pp. 159–170.

44. Macalalad, A. R., Zody, M. C., Charlebois, P., Lennon, N. J., Newman, R. M., Malboeuf, C. M., Ryan, E. M., Boutwell, C. L., Power, K. A., Brackney, D. E., et al. (2012) Highly sensitive and specific detection of rare variants in mixed viral populations from massively parallel sequence data. PLoS computational biology, 8(3), e1002417.

45. Yang, X., Charlebois, P., Macalalad, A., Henn, M. R., and Zody, M. C. (2013) V-Phaser 2: variant inference for viral populations. BMC genomics, 14(1), 674.

46. Routh, A., Chang, M. W., Okulicz, J. F., Johnson, J. E., and Torbett, B. E. (2015) CoVaMa: Co-Variation Mapper for disequilibrium analysis of mutant loci in viral populations using next-generation sequence data. Methods, 91, 40–47.

47. Verbist, B. M., Thys, K., Reumers, J., Wetzels, Y., Van der Borght, K., Talloen, W., Aerssens, J., Clement, L., and Thas, O. (2014) VirVarSeq: a low-frequency virus variant detection pipeline for Illumina sequencing using adaptive base-calling accuracy filtering. Bioinformatics, 31(1), 94–101.

48. Artyomenko, A., Wu, N. C., Mangul, S., Eskin, E., Sun, R., and Zelikovsky, A. (June, 2017) Long Single-Molecule Reads Can Resolve the Complexity of the Influenza Virus Composed of Rare, Closely Related Mutant Variants. Journal of Computational Biology, 24(6), 558–570.

49. Mangul, S., Martin, L. S., Hill, B. L., Lam, A. K.-M., Distler, M. G., Zelikovsky, A., Eskin, E., and Flint, J. (March, 2019) Systematic benchmarking of omics computational tools. Nature Communications, 10(1).

50. Mitchell, K., Brito, J. J., Mandric, I., Wu, Q., Knyazev, S., Chang, S., Martin, L. S., Karlsberg, A., Gerasimov, E., Littman, R., Hill, B. L., Wu, N. C., Yang, H. T., Hsieh, K., Chen, L., Littman, E., Shabani, T., Enik, G., Yao, D., Sun, R., Schroeder, J., Eskin, E., Zelikovsky, A., Skums, P., Pop, M., and Mangul, S. (March, 2020) Benchmarking of computational error-correction methods for next-generation sequencing data. Genome Biology, 21(1).

51. Eliseev, A., Gibson, K. M., Avdeyev, P., Novik, D., Bendall, M. L., Pérez-Losada, M., Alexeev, N., and Crandall, K. A. (August, 2020) Evaluation of haplotype callers for next-generation sequencing of viruses. Infection, Genetics and Evolution, 82, 104277.

52. Giallonardo, F. D., Töpfer, A., Rey, M., Prabhakaran, S., Duport, Y., Leemann, C., Schmutz, S., Campbell, N. K., Joos, B., Lecca, M. R., Patrignani, A., Däumer, M., Beisel, C., Rusert, P., Trkola, A., Günthard, H. F., Roth, V., Beerenwinkel, N., and Metzner, K. J. (2014) Full-length haplotype reconstruction to infer the structure of heterogeneous virus populations. Nucleic Acids Research, 42(14), e115.

53. Zanini, F., Brodin, J., Thebo, L., Lanz, C., Bratt, G., Albert, J., and Neher, R. A. (Dec, 2015) Population genomics of intrapatient HIV-1 evolution. eLife,.

54. Benidt, S. and Nettleton, D. (2015) SimSeq: a nonparametric approach to simulation of RNA-sequence datasets. Bioinformatics, 31(13), 2131–2140.

55. Gerasimov, E. Analysis of NGS Data from Immune Response and Viral Samples PhD thesis Georgia State University (2017).

56. Levina, E. and Bickel, P. (2001) The EarthMover’s Distance is the Mallows Distance: Some Insights from Statistics. Proceedings of ICCV, 2001 pp. 251–256.

57. Mallows, C. L. (1972) A note on asymptotic joint normality. Annals of Mathematical Statistics, 43(2), 508–515.

58. Yang, X., Charlebois, P., Gnerre, S., Coole, M. G., Lennon, N. J., Levin, J. Z., Qu, J., Ryan, E. M., Zody, M. C., and Henn, M. R. (2012) De novo assembly of highly diverse viral populations. BMC genomics, 13(1), 475.

59. Artyomenko, A., Wu, N. C., Mangul, S., Eskin, E., Sun, R., and Zelikovsky, A. (2017) Long Single-Molecule Reads Can Resolve the Complexity of the Influenza Virus Composed of Rare, Closely Related Mutant Variants. Journal of Computational Biology, 24(6), 558–570.

60. Bron, C. and Kerbosch, J. (September, 1973) Algorithm 457: Finding All Cliques of an Undirected Graph. Commun. ACM, 16(9), 575–577.

61. Kováč, J. (July, 2013) Complexity of the Path Avoiding Forbidden Pairs Problem Revisited. Discrete Appl. Math., 161(10-11), 1506–1512.

62. Nicolae, M., Mangul, S., Mandoiu, I., and Zelikovsky, A. (2011) Estimation of alternative splicing isoform frequencies from RNA-Seq data. Algorithms for Molecular Biology, 6:9.

63. Kireev, D. E., Lopatukhin, A. E., Murzakova, A. V., Pimkina, E. V., Speranskaya, A. S., Neverov, A. D., Fedonin, G. G., Fantin, Y. S., and Shipulin, G. A. (11, 2018) Evaluating the accuracy and sensitivity of detecting minority HIV-1 populations by Illumina next-generation sequencing. J. Virol. Methods, 261, 40–45.

64. Wertheim, J. O., Leigh Brown, A. J., Hepler, N. L., Mehta, S. R., Richman, D. D., Smith, D. M., and Kosakovsky Pond, S. L. (2014) The Global Transmission Network of HIV-1. The Journal of Infectious Diseases, 209(2), 304–313.

65. Quail, M. A., Smith, M., Coupland, P., Otto, T. D., Harris, S. R., Connor, T. R., Bertoni, A., Swerdlow, H. P., and Gu, Y. (2012) A tale of three next generation sequencing platforms: comparison of Ion Torrent, Pacific Biosciences and Illumina MiSeq sequencers. BMC genomics, 13(1), 341.

