## Supplementary Material for "Accurate assembly of minority viral haplotypes from next-generation sequencing through efficient noise reduction"

### Supplementary Results

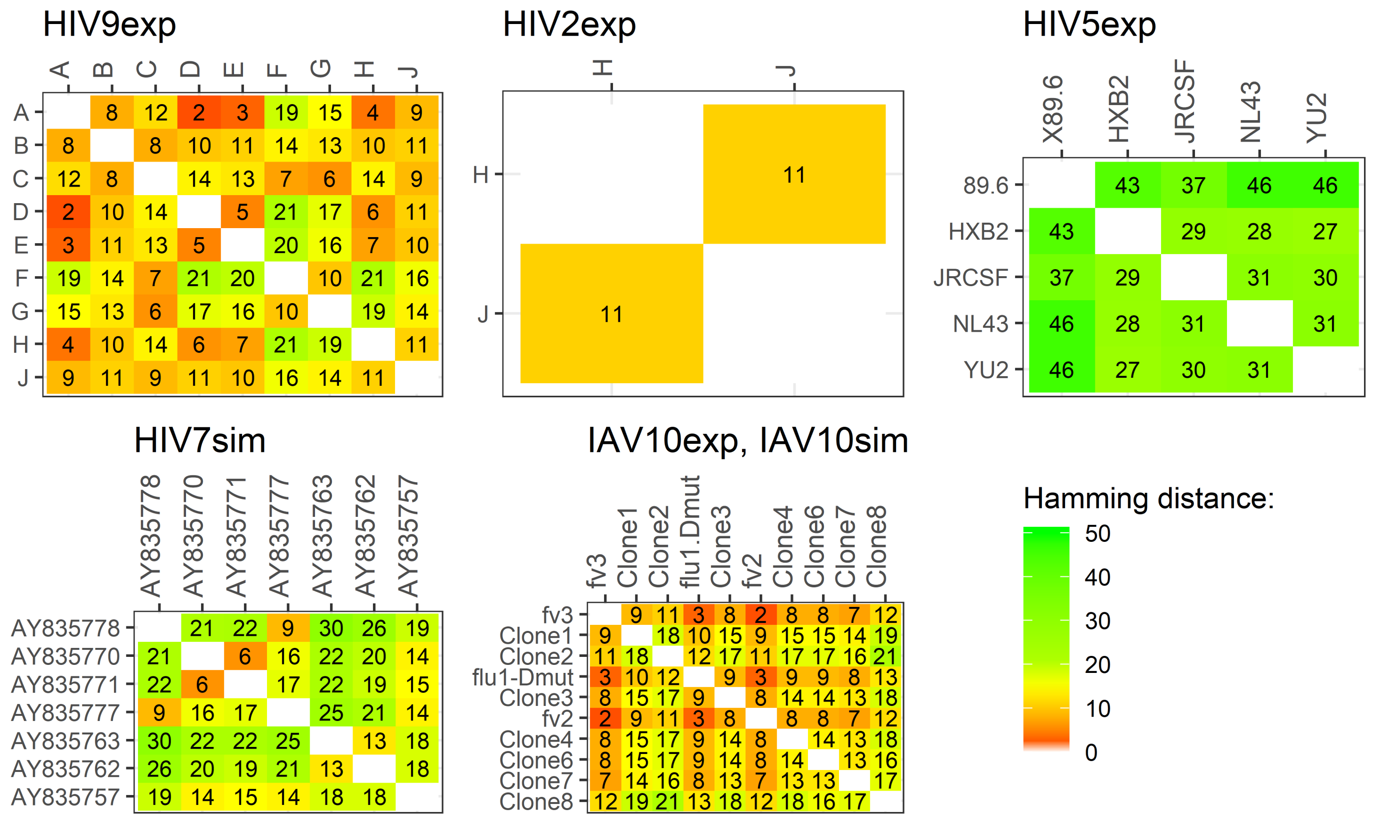


**Figure S1.** Pairwise Hamming distances between variants in the experimental (exp) and simulated (sim) datasets HIV9exp, HIV2exp, HIV5exp, HIV7sim, IAV10sim, and IAV10exp.

| PacBio # of Reads | Method | Variant | fv3 | Clone1 | Clone2 | flu1-Dmut | Clone3 | fv2 | Clone4 | Clone5 | Clone6 | Clone7 | FP |
|---|---|---|---|---|---|---|---|---|---|---|---|---|---|
| | | True Freq.,% | 50 | 25 | 12.5 | 6.25 | 3.125 | 1.56 | 0.78 | 0.39 | 0.19 | 0.097 | |
| 33.5K (all) | CliqueSNV | Match | ✓ | ✓ | ✓ | ✓ | ✓ | ✓ | ✓ | ✓ | ✓ | ✓ | 0 |
| | | Freq., % | 52.6 | 23.7 | 12.6 | 6.4 | 2.3 | 1.17 | 0.7 | 0.35 | 0.12 | 0.051 | 0 |
| | 2SNV | Match | ✓ | ✓ | ✓ | ✓ | ✓ | ✓ | ✓ | ✓ | ✓ | × | 1 |
| | | Freq., % | 51.8 | 23.7 | 12.5 | 6.4 | 2.3 | 1.2 | 0.7 | 0.3 | 0.1 | 0 | 1.0 |
| | PredictHaplo | Match | ✓ | ✓ | ✓ | × | ✓ | × | ✓ | ✓ | × | × | 0 |
| | | Freq.,% | 56.7 | 23.8 | 13.7 | 0 | 3.1 | 0 | 1.5 | 1.2 | 0 | 0 | 0 |
| Subsampling | | | | | | | | | | | | | |
| 16K | CliqueSNV | Match,% | 100 | 100 | 100 | 100 | 100 | 90 | 100 | 100 | 100 | 20 | 0.1 |
| | | Freq., % | 52.9 | 23.7 | 12.5 | 6.4 | 2.3 | 1.19 | 0.71 | 0.32 | 0.12 | 0.69 | 1.15 |
| | 2SNV | Match,% | 100 | 100 | 100 | 100 | 100 | 100 | 100 | 100 | 0 | 0 | 0.2 |
| | | Freq., % | 52.4 | 23.7 | 12.5 | 6.4 | 2.3 | 1.1 | 0.7 | 0.3 | 0 | 0 | 0.6 |
| | PredictHaplo | Match | 100 | 100 | 100 | 70 | 100 | 0 | 100 | 40 | 0 | 0 | 0.3 |
| | | Freq.,% | 54.2 | 23.5 | 13.1 | 6.0 | 2.9 | 0 | 1.4 | 1.0 | 0 | 0 | 0.5 |
| 8K | CliqueSNV | Match,% | 100 | 100 | 100 | 100 | 100 | 90 | 100 | 100 | 30 | 0 | 0 |
| | | Freq., % | 52.8 | 23.6 | 12.5 | 6.5 | 2.3 | 1.2 | 0.7 | 0.35 | 0.16 | 0 | 0 |
| | 2SNV | Match,% | 100 | 100 | 100 | 100 | 100 | 100 | 100 | 0 | 0 | 0 | 0 |
| | | Freq., % | 53.1 | 23.7 | 12.5 | 6.5 | 2.3 | 1.25 | 0.7 | 0 | 0 | 0 | 0 |
| | PredictHaplo | Match,% | 100 | 100 | 100 | 0 | 100 | 0 | 100 | 20 | 0 | 0 | 0.2 |
| | | Freq.,% | 58.1 | 24.0 | 12.7 | 0 | 3.1 | 0 | 1.6 | 1.3 | 0 | 0 | 0.5 |
| 4K | CliqueSNV | Match,% | 100 | 100 | 100 | 100 | 100 | 80 | 100 | 40 | 0 | 0 | 0 |
| | | Freq., % | 53.3 | 23.7 | 12.3 | 6.4 | 2.4 | 1.19 | 0.7 | 0.39 | 0 | 0 | 0 |
| | 2SNV | Match,% | 100 | 100 | 100 | 100 | 100 | 100 | 20 | 0 | 0 | 0 | 0 |
| | | Freq., % | 53.7 | 23.7 | 12.3 | 6.5 | 2.4 | 1.2 | 0.9 | 0 | 0 | 0 | 0 |
| | PredictHaplo | Match,% | 100 | 100 | 100 | 0 | 70 | 0 | 10 | 0 | 0 | 0 | 0.3 |
| | | Freq.,% | 60.1 | 23.9 | 12.8 | 0 | 3.5 | 0 | 2.5 | 0 | 0 | 0 | 0.5 |

**Table S1.** Comparison of CliqueSNV, 2SNV and PredictHaplo on full and sub-sampled data ***(PacBio, experimental)***. For all 33.5K reads, the sign "✓" (respectively, "×") denotes fully matched (respectively, unmatched) true variant and the column FP reports the number of incorrectly predicted variants (false positives) and their total frequency. For each sub-sample size (16K,...,4K), the table reports the percent of runs when a variant is completely matched and its average frequency over runs when the variant was detected. Similarly, the column FP reports the average number of false positive variants and their average total frequency. Colors indicate the percent of matched variants: green - high percent, red - low percent.

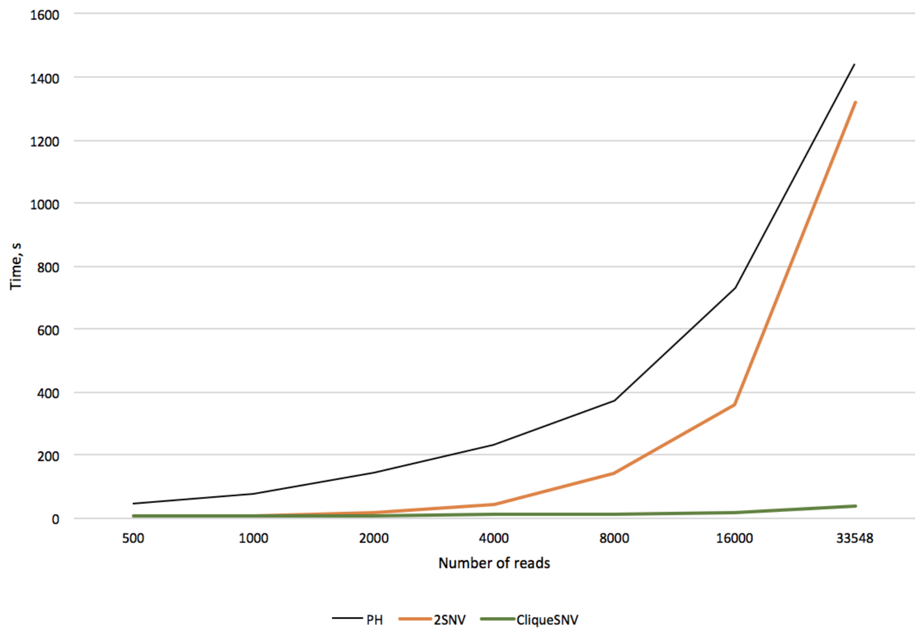


**Figure S2.** Runtimes of PredictHaplo (PH), 2SNV and CliqueSNV on datasets with different read sizes.

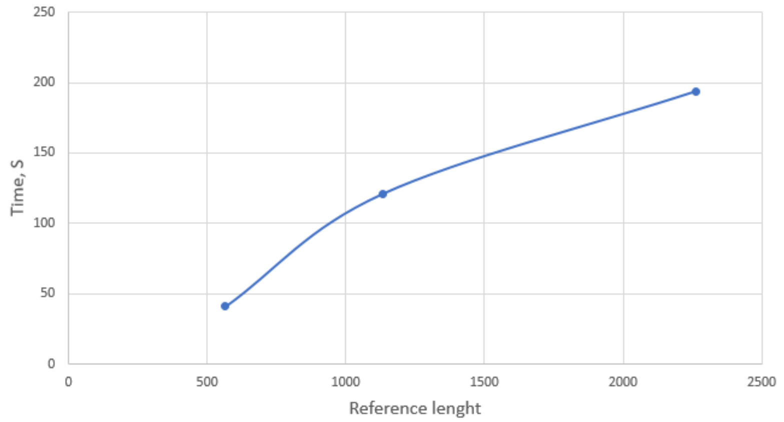


**Figure S3.** CliqueSNV runtime on datasets with different reference length and same coverage (about 1M reads in total).

### Supplementary Methods

#### *Benchmark preparation*

We used 50,000 total copies of plasmid DNA from these nine constructs as input for a nested PCR reaction to amplify the polymerase region using the following primary and nested primers respectively: HIV-B PRO-OUT.3F, 5' CCT CAG ATC ACT CTT TGG CAA CG 3' and HIV-RT 215/219.3R, 5' CTT CTG TAT GTC ATT GAC AGT CC 3' Nested PCR: HIV-B PR/RT.2F, 5' CTT TGG CAA CGA CCC CTY GTC CA 3' and HIV-RT 181-190.1.4R, 5' ATC AGG ATG GAG TTC ATA ACC CA 3'.

The primary and nested PCRs were done using 94°C for four minutes, followed by 40 cycles of 94°C for one minute, 50°C for 30 seconds, and 72°C for two minutes and a final extension at 72°C for five minutes[1].

We created two plasmid mixtures to generate artificial mixtures simulating clinical specimens containing many variants at different virus levels. The mixtures comprised nine and two plasmids with varying copy numbers of each plasmid.

PCR reactions were generated and purified using the QIAquick PCR purification kit. (Qiagen, Valencia CA) The purified amplicons (10 ng) were subsequently used for NGS library construction using the Nextera XT DNA Library Prep kit (Illumina Inc., San Diego, CA). Libraries were pooled, and enriched for 900-1,000-bp fragments using magnetic bead based size selection (AMPure XP, Beckman Coulter, Brea, CA) and sequenced on a MiSeq v3 (600-cycle) flow cell on the Miseq system ( Illumina Inc., San Diego CA).

#### *Pseudocode of the CliqueSNV algorithm*

**Algorithm 1** CliqueSNV Algorithm

**Step 1: finding linked and forbidden SNV pairs**
- Split the read alignment $M_{L\times N}$ into binary matrix $4M$
- Construct a compact representation of the binary matrix $4M$
- For each $I,J \in \{1,\dots,4L\}$ find $O^{IJ}$ and $O^{IJ}_{22}$, where
  - $O^{IJ}$ = # of reads covering both $I$ and $J$
  - $O^{IJ}_{22}$ = # of reads with both minor SNVs
  - If $O^{IJ}_{22} > \varepsilon O^{IJ}$ compute $p$-value (default $\varepsilon = 0.0003$)
- Find all linked SNV pairs with the adjusted p-value $< 1\%$

**Step 2: constructing the SNV graph**
- Filter out 10% of the most erroneous PacBio reads
- Construct the SNV graph $G = (V,E)$, where
  - $V = \{1,\dots,4L\}$, and $E$ are links between minor SNVs

**Step 3: finding maximal cliques in the SNV graph using Bron-Kerbosch algorithm[2]**

**Step 4: merging cliques in the clique graph with forbidden pairs**
- Find the clique graph $C_G$ with pairs.
- Find all maximal connected subgraphs in $C_G$.
- Merge all cliques inside each maximal connected subgraph.

**Step 5: partitioning reads between merged cliques and finding consensus haplotypes**
- Find the set $S$ of all positions that belong to at least one clique.
- Make an empty clique on $S$.
- Assign each read to the closest clique.
- Find the consensus $v(q)$ of all assigned reads for each $q$.

**Step 6: estimating haplotype frequencies by expectation-maximization algorithm**

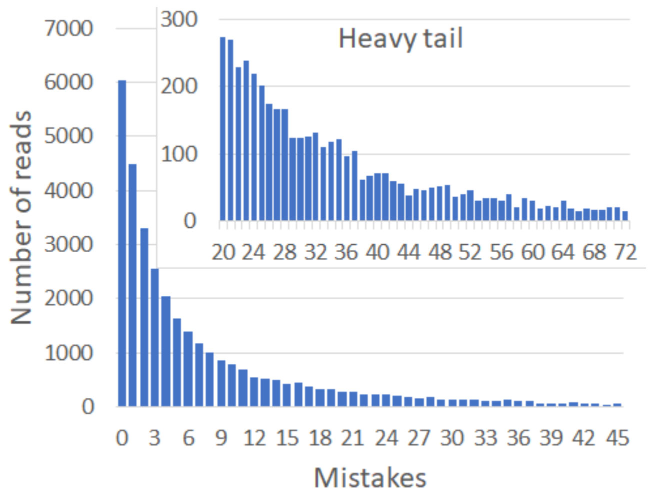


**Figure S4.** A typical distribution of errors in PacBio reads. The heavy tail indicates that a significant portion of errors is accumulated by a relatively small number of reads.

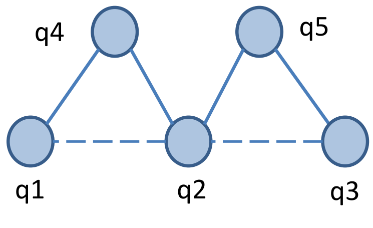


**Figure S5.** The clique graph $C_G$ with 5 vertice corresponding to cliques in $G$, 4 edges and two forbidden pairs $(q_1,q_2)$ and $(q_2,q_3)$. There 3 maximal connected subgraphs avoiding forbidden pairs: $\{q_1,q_4\}$ $\{q_4,q_2,q_5\}$ $\{q_5,q_3\}$

### References


**1.** Peters, P. J. *et al.* Hiv infection linked to injection use of oxymorphone in indiana, 2014–2015. *New Engl. J. Medicine* **375**, 229–239 (2016).

**2.** Bron, C. & Kerbosch, J. Algorithm 457: Finding all cliques of an undirected graph. *Commun. ACM* **16**, 575–577, DOI: 10.1145/362342.362367 (1973).
